# Strong gene activation with genome-wide specificity using a new orthogonal CRISPR/Cas9-based Programmable Transcriptional Activator

**DOI:** 10.1101/486068

**Authors:** S Selma, J Bernabé-Orts, M Vazquez-Vilar, B Diego, M Ajenjo, V García-Carpintero, A Granell, D Orzaez

**Affiliations:** Instituto de Biología Molecular y Celular de Plantas (IBMCP), Consejo Superior de Investigaciones Científicas, Universidad Politécnica de Valencia, Camino Vera s/n, 46022 Valencia, Spain.

## Abstract

Synthetic Biology (SynBio) aims at rewiring plant metabolic and developmental programs with orthogonal regulatory circuits. This endeavour requires new molecular tools able to interact with endogenous factors in a potent yet at the same time highly specific manner. A promising new class of SynBio tools that could play this function are the synthetic transcriptional activators based on CRISPR/Cas9 architecture, which combine autonomous activation domains (ADs) capable of recruiting the cell’s transcription machinery, with the easily customizable DNA-binding activity of nuclease-inactivated Cas9 protein (dCas9), creating so-called Programmable Transcriptional Activators (PTAs). In search for optimized dCas9-PTAs we performed a combinatorial analysis with seven different ADs arranged in four different protein/RNA architectures. This analysis resulted in the selection of a new dCas9-PTA with improved features as compared with previously reported activators. The new synthetic riboprotein, named dCasEV2.1, combines EDLL and VPR ADs using a multiplexable mutated version (v2.1) of the previously described aptamer-containing guide RNA2.0. We show here that dCasEV2.1 is a strong and wide spectrum activator, displaying variable activation levels depending on the basal activity of the target promoter. Maximum activation rates reaching up to 10000 fold were observed when targeting the NbDFR gene. Most remarkably, RNAseq analysis of dCasEV2.1-transformed *N. benthamiana* leaves revealed that the topmost activation capacity of dCasEV2.1 on target genes is accompanied with strict genome-wide specificity, making dCasEV2.1 an attractive tool for rewiring plant metabolism and regulatory networks.

## INTRODUCTION

The ability to superimpose synthetic regulatory circuits on plant endogenous gene expression networks has been aimed for a long time as it will open a range of applications in plant biotechnology. Classical attempts have involved the ectopic expression of heterologous transcriptional factors (TFs) under the control of purpose-specific promoters, therefore connecting promoter-specified inputs to a cascade of TF-targeted activated/repressed genes as output ^1^. An important limitation of this approach is that does not allow free selection of the output response, as the collection of target genes is restricted by the DNA binding specificities of the TFs employed, which are typically “hardwired” at the protein level. A way to circumvent this limitation is to engineer artificial promoters with TF-binding regulatory boxes controlling all target genes ^2–4^. However, this requires important engineering efforts, limiting its applicability in practical terms.

Furthermore, “natural” TFs have often a broad spectrum of DNA binding activities, limiting the orthogonality and specificity of the approach.

Contrary to hardwired TFs, CRISPR/Cas9 protein architecture enables the design of transcription factors with DNA-binding specificities can be soft-programmed in a 20 nucleotide-long guide RNA with minimum engineering efforts. Using catalytically inactive Cas9 (dCas9) nuclease fused to transcriptional activator or repressor domains, the gene expression of an individual gene can be modified ^5^. The CRISPR-dCas9 strategy presents advantages compared with previously described modifiable regulators including zinc finger nucleases ZFNs ^6,7^ or TAL effectors ^8,9^ which are very specific but require a new recoding of regulatory protein for each target sequence.

Initial attempts to produce CRISPR-dCas9 programmable transcriptional activators (Cas9-PTAs) in plants made use of transcriptional activator domains (TAD) fused to dCas9 protein, achieving only low/moderate activation rates ^10,11^. Next generation of PTAs are designed to achieve increased activation rates by combining several TADs displayed in a single dCas9 protein. To do so, different strategies have been proposed. The SunTag strategy ^12^ uses multi-epitope tags to attach multiple TADs, whereas SAM and scRNA strategies ^13–15^ use RNA aptamers added to the gRNA scaffold as secondary anchoring sites for TADs. All those strategies have been showed to improve activation rates in mammalian cells, and recently shown similar results in plants ^16,17^. Despite these achievements, the search for optimal PTAs in plants is still open to improved designs. Ideally, plant PTAs should combine strong activation rates with a wide spectrum of responsive targets and most importantly, with high target specificity, that is, ensuring that the cell transcriptome is only affected in the intended gene(s).

In a search for improved plant dCas9-PTAs, here we show the results of a systematic comparison of 43 SunTag, SAM and scRNA-based TAD combinations tested for their ability to activate different promoters fused to a Luciferase reporter. As a result of these analysis, we selected a new dCas9-PTA comprising two TADs (EDLL and VPR) next to an aptamer variant of the gRNA2.0 scaffold employed in the scRNA strategy. The new dCas9-PTA (named as dCasEV2.1) consistently produced the highest activation rates in all assays, both using transiently-transformed and genome-integrated promoters as targets. dCasEV2.1 was able to activate the *Nicotiana benthamiana* Dihydroflavonol-4-reductase (DFR), an inducible gene with very low basal activity levels, up to 10000 folds. When directed to strong constitutive promoters, dCasEV2.1 yielded lower induction rates, but raised transcriptional activity up to levels that triplicate those of the strong CaMV35S promoter. Moreover, RNAseq analysis showed a remarkable genome-wide specificity of dCasEV2.1-PTA, with virtually no changes in the transcriptome other than those anticipated by off-target analysis.

## RESULTS

A first round of comparisons among the different PTA dCas9-strategies was performed transiently in *N. benthamiana* leaves. PTA transactivation levels were assessed using Nopaline synthase promoter (pNos) coupled to firefly luciferase (Fluc) reporter. A constitutive Renilla luciferase (Rluc) was used as internal reference. For initial comparisons, PTAs were targeted to the pNos promoter with a single gRNA annealing at position -161 relative to the Transcriptional Start Site (TSS) (Fig 1A). This position was validated in previous experiments ^10^. SAM, scRNA, and SunTag (Fig 1B) designs were analysed using the activation domains VP64 and EDLL. For SunTag, VP64 and EDLL were fused to the ScFv, and tested separately. For scRNA and SAM, EDLL was attached to dCas9 and VP64 was fused to the MS2 viral coat protein, which binds an RNA aptamer. RNA scaffolds in scRNA and SAM designs contained a second optimized aptamer next to the wild type one, as this double-aptamer design was earlier found to improve binding activity ^18^. During the cloning of the gRNA2.0 scaffold employed in the scRNA design, a spontaneous mutation occurred consisting in the insertion of an adenine in the loop of the first aptamer (Supplementary Figure 2). Since this aptamer variant had not been studied earlier, we decided to include it in the comparison analysis (labelled as gRNA2.1). The experiment was completed with direct dCas9:VP64 and dCas9:EDLL fusions, used to monitor the improvement achieved from earlier designs. Relative transcriptional activities (RTAs) in transient assays were expressed as relative transcription units (RPUs). RPUs are calculated as the Fluc/Rluc ratios measured in each sample, normalized with the Fluc/Rluc ratios produced by an un-activated pNos promoter (GB1398) assayed in parallel. This procedure was earlier proposed as a standard measurement for the documentation of standard DNA parts (phytobricks)^19,20^. In these conditions, the CaMV35S promoter-based standard DNA part GB0164 consistently confers activity levels of 12 +/-2 RPUs. Thus, this CaMV35S reporter was included in all comparative experiments to serve as an upper limit reference.

**Figure 1.**
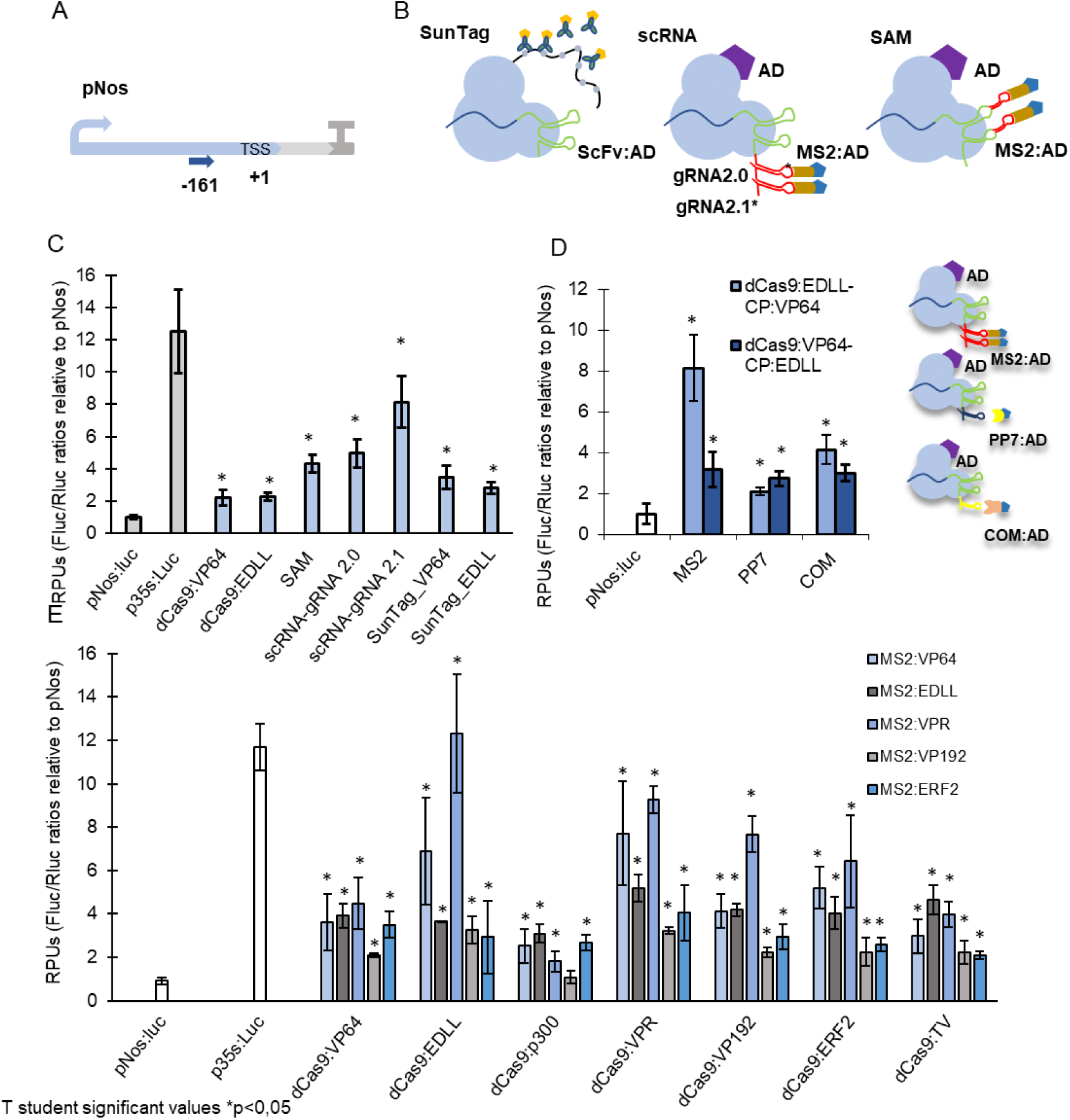
Transient activation of pNos promotor by dCas9-PTAs. (A) Representation of pNos promoter with a gRNA target at position -161. (B) Schematic representation of SunTag-strategy, scRNA-strategy (with gRNA2.0 or gRNA 2.1 alternative scaffolds), and SAM-strategy used in activation experiments. Asterisk indicates the mutation position in gRNA2.1 (C) Relative transcriptional activities (RTAs) obtained with different dCas9-PTAs strategies: dCas9:VP64 and dCas9:EDLL represent direct TAD fusion strategies; All SAM, scRNA and SunTag combinations follow the architecture represented in (A). All SAM and scRNA strategies combine direct Cas9-EDLL fusions with gRNA aptamers attached to MS2-VP64. (D) RTAs obtained upon activation of pNos at -161 position with different aptamer-coat protein combinations using scRNA architecture. MS2 strategy combines gRNA2.1 with MS2 coat protein as in (A); PP7 and COM strategies use a single aptamer to bind PP7 and COM coat proteins respectively; coloured bars represent the two different TAD combinations assayed in each strategy. Right panel shows a schematic representation of the three strategies. (E) RTAs obtained upon activation with different combination of TADs using scRNA-gRNA 2.1 strategy and targeting the reporter pNos in position -161. pNos:Luc is a non-activated pNos promoter Fluc/Rluc reporter; p35S:luc is a Fluc/Rluc reporter driven by a standard 35S promoter. RTAs are measured as relative transcriptional units (RPUs) calculated as the Fluc/Rluc ratios for each construct normalized with the Fluc/Rluc ratio of a pNos:Luc reporter transformed in parallel. Bars represent average RTAs +/- SD, n=3. Asterisks indicate T student significant values (p<0,05)

As shown in Fig 1C, scRNA, SAM and SunTag strategies produced higher activation rates as compared with previous simplified designs of EDLL or VP64 dCas9 fusions. Furthermore, designs involving combinations of different TADs (scRNA and SAM) showed stronger transactivation than the SunTag design, which involves multiple copies of the same TAD. Surprisingly, the highest transcriptional activation was achieved with scRNA-gRNA2.1 scaffold, reaching RTA levels close to CaMV35S (RTA= 8 +/-2 RPUs) and outperforming previously optimized scRNA-gRNA2.0 strategy. In a separate experiment, the gRNA2.1 was compared with similar aptamer-binding coat proteins (CP) PP7 and COM ^15^ using the equivalent TAD combinations dCas9:EDLL-CP:VP64 and dCas9:VP64-CP:EDLL (Fig 1D). Again, the gRNA2.1 showed the best activation results. Note that by swapping the position of the activation domains, different levels of activation were achieved.

In a further optimization step, and attending to the differences obtained in TAD swapping experiments described above, we tested the scRNA2.1 design with new TAD combinations as shown in Fig 1E. Seven previously described TADs (VP64, EDLL, P300, VPR, VP192, ERF2 and TV) were fused to dCas9 and combined with five of them (VP64, EDLL, VPR, VP192 and ERF2) anchored at the MS2-aptamer position. A detailed description for each of the TADs employed in this experiment is shown in Table 1. VPR domain showed the highest activation rates in most combinations assayed, followed by VP64 and EDLL. Remarkably, the dCas9:EDLL-MS2:VPR gave maximum pNos activation up to RTA levels similar to the CaMV35S GB0164 used as our upper limit reference. A separate experiment shown in additional figure (Supplementary Figure 3B) confirmed that, as expected, the selected dCas9:EDLL-MS2:VPR tandem performed better when bound to the newly described gRNA2.1 aptamer.

**Table 1.** Description of TADs employed in combinatorial assays.

| Transcriptional Activation Domain (TAD) | Description | Reference |
| --- | --- | --- |
| VP64 | Tetrameric repeat of herpes simplex VP16. | <sup>36</sup> |
| EDLL | Short motif present in *Arabidopsis* AP2 sub-family. Activation domain. | <sup>37</sup> |
| P300 | Human acetylase of histone H3 lysine 27 P300 core | 38 |
| VPR | (VP64-p65-Rta) Tripartite transactivation domain. p65 is a subunit from NF-kappa B transcription factor that contains the trans-activation domain. Rta is Epstein-Barr virus R transactivator. | 39 |
| VP192 | Trimeric repeat of VP64 | 40 |
| ERF2(m) | Modified trans-activation domain of *Arabidopsis* ERF2 protein involved in ethylene response. | 41 |
| TV | 6xTAL - VP128. Six copies of TALE Trans-activation domain motif from *Xanthomonas* and dimeric repeat of VP64. | 21 |

The results obtained with the pNos:Luc reporter prompted us to test the activity of dCas9:EDLL-MS2:VPR/gRNA2.1 (now abbreviated as dCasEV2.1) on promoters with contrasting basal activities. The promoter derived from the tomato SlMTB (Metallothionein-like protein type 2 B) gene, (catalogued as GB1399) has a strong constitutive basal activity (RTA= 4 +/-1 RPUs) and could serve to test dCasEV2.1 ability to rise transcription levels in absolute terms. On the contrary, the promoter of the SlDFR gene (Dihydroflavonol-4-reductase) from *Solanum licopersicum* (pSlDFR, catalogued as GB1160), has very low basal expression levels (RTA < 0.04 RPUs), but it is strongly induced *in planta* by the presence of “natural” Myb TFs (e.g. SlANT1). pSlDFR was used as model promoter to test the activation range of dCasEV2.1 induction.

Activation of the pSlMTB was first analysed using dCasEV2.1 complex and gRNAs at positions -98, -129, -184 and -541, represented in Fig 2A. The gRNAs were tested individually or combined in a single T-DNA with the GoldenBraid multiplexing cloning strategy. As observed (Fig 2B), all gRNA tested in a range of 500 bps upstream the TSS conferred strong activation to the promoter, with position -129 reaching maximum levels. The combination of gRNAs, in this case, did not confer activation levels higher than those obtained by gRNAs acting individually. Most notably, absolute RTA levels obtained with gRNA -129 reached record RTA levels (60+/-10 RPUs), corresponding to a 20 fold activation from pMTB basal levels and 4 times above CaMV35 levels used as upper limit reference. The superior performance of the dCasEV2.1 complex was once more confirmed in a separate activation experiment with gRNAs in position -98, -129, -184 and -541 in combination and a selected number of alternative gRNA2.1 combinations (dCas9:VPR-Ms2:VPR and dCas9:TV-Ms2:VPR). As shown (Fig 2C) the results confirm that dCasEV2.1 achieved the best activation rates also when using pSlMTB as target promoter.

**Figure 2.**
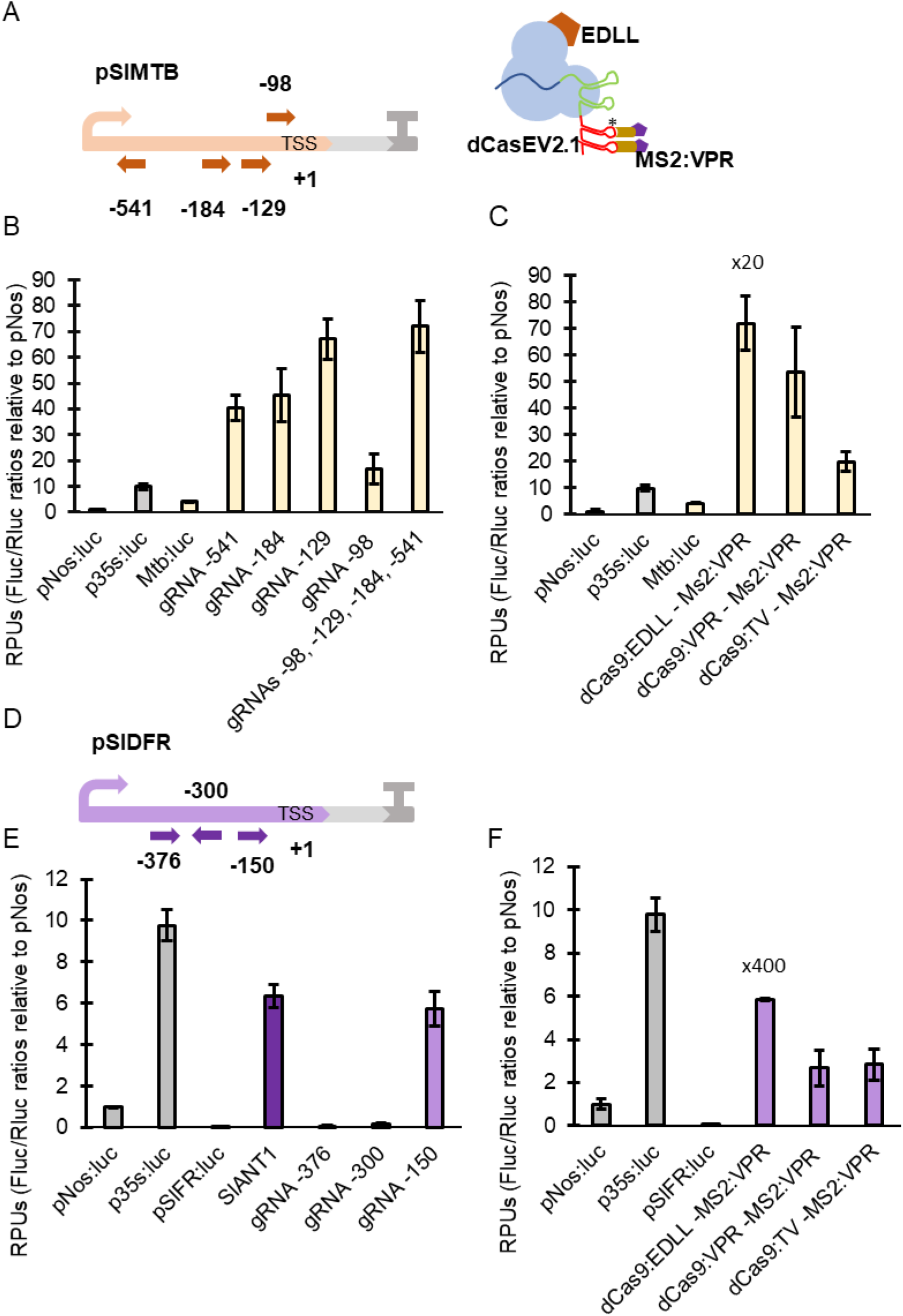
Transient activation of SlMTB and SlDFR promoter by dCas9-PTAs. (A) Representation of the reporter pSlMTB with gRNAs targeted at positions -98, -129, -184 and -541 with dCasEV2.1 (B) Relative transcriptional activities (RTAs) obtained with gRNAs targeting reporter pSlMTB at positions -98, -129, -184, and -541, individually and in combination, using dCasEV2.1. (C) RTAs obtained upon activation with the best combination of TADs (dCas9:EDLL-MS2:VPR, dCas9:VPR-MS2:VPR and dCas9:TV-MS2:VPR) using scRNA-gRNA 2.1 strategy and targeting the reporter pSlMTB at positions -98, -129, -184, and -541. (D) Representation of the reporter pSlDFR with gRNAs targeted at positions -150, -300, and -376. (E) Relative transcriptional activities (RTAs) obtained with gRNAs targeting reporter SlDFR at positions -150, -300, and -376 individually, using dCasEV2.1. In this experiment Myb-TF SlANT1 was included to test the RTA obtained with the natural transcription factor of SlDFR. (F) RTAs obtained upon activation with the best combinations of TADs (dCas9:EDLL-MS2:VPR, dCas9:VPR-MS2:VPR and dCas9:TV-MS2:VPR) using scRNA-gRNA 2.1 strategy and targeting the reporter pSlDFR at position -150. RTAs are measured as relative promoter units (RPUs) calculated as the Fluc/Rluc ratios for each construct normalized with the Fluc/Rluc ratio of a pNos:Luc reporter transformed in parallel. Bars represent average RTAs +/- SD, n=3.

Next, the activation of SlDFR promoter by Cas9-PTAs was analysed. The SlDFR gene is involved in flavonoids biosynthesis pathway and it is strongly induced by natural transcription factors of the MYB family. Three gRNAs targeting the positions -376, -300 and -150 were generated and tested with scRNA2.1. The pSlDFR activation assay was also interrogated with SlANT1, a Myb factor than naturally activates the DFR gene in tomato. Best activation rates were obtained with gRNA that targets at position -150 (×100 fold) (Fig 2E). As in previous promoters assayed, dCasEV2.1 outperformed other AD combinations also in the case of pSlDFR (Fig 2F).

Transient experiments with Agrobacterium-delivered T-DNAs allow fast combinatorial dCas9-PTA transactivation assays, however the episomal status of at least part of the transcriptionally active T-DNAs used as reporters may lead to regulatory dynamics different to those of chromatin-imbibed genes, which are often influenced by nucleosome position and subjected to epigenetic regulation. Therefore, we next studied dCasEV2.1 activity on chromatin-integrated genes, using two type of strategies: (i) by targeting reporter promoters integrated in transgenic lines and (ii) by assessing activation endogenous *N. benthamiana* genes.

Transgenic reporter lines were generated by transforming *N. benthamiana* with T-DNA constructs used in transient experiments plus the addition of a KanR module. Single-copy T2 lines Nos-RL3, Nos-RL5 and Nos-RL6 (for pNos promoter), MTB-RL3 (for pMTB promoter) and DfrRL1 (for pDFR promoter) were selected for activation tests after analysing their KanR segregation. For each line, leaves were agroinfiltrated using dCasEV2.1 activating construct with or without target-specific gRNAs, and the activation levels measured as Luc/Ren ratios. As shown in Fig 3A, activation rates were in line with what was observed in transient experiments. Chromatin-integrated pNos was induced 3- 13 fold reaching maximum levels similar to CaMV35S promoter; the pMTB line, showing higher basal levels, was only activated 3X, but reached record expression levels in absolute terms (as compared with a agroinfiltrated CaMV35S promotor used as reference). DFR-RL1 line having lowest basal luminescence rates, showed highest activation rate up to 90 fold.

**Figure 3.**
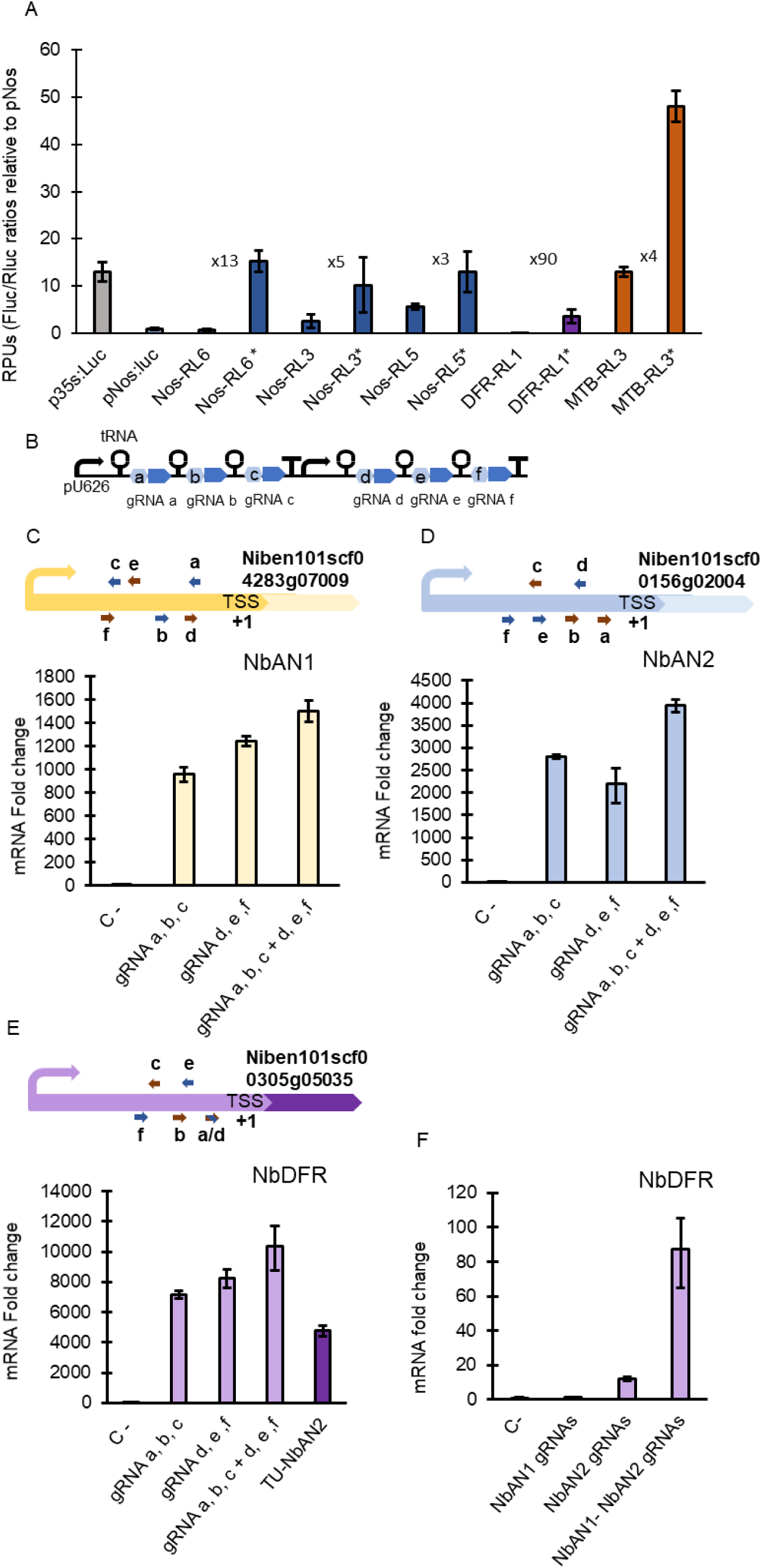
Transient activation of transgenic reporter lines and endogenous genes *in N. benthamiana* with dCasEV2.1. (A) Relative transcriptional activities (RTAs) obtained of the activation of Nos, SlDFR and MTB reporter transgenic lines using dCasEV2.1. The Nos-RL6, Nos-RL3 and Nos-RL5 were activated targeting the pNos reporter at positions -161 and -211. DFR-RL1 was activated targeting the SlDFR promoter at position -150. MTB-RL3 was activated targeting the SlMTB promoter at positions -98, -129, -184 and -541. Asterisks represent the reporter line activation using dCasEV2.1 (B) Representation of the Multiplexing gRNA strategy that adds the scRNA-gRNA 2.1. (C) mRNA fold change obtained targeting the endogenous gene NbAN1 with dCasEV2.1 through the multiplexing gRNA strategy after 4 dpi. Represented gRNAs target the promoter of NbAN1 in the positions: a (-101), b (-173), c (-242), d (-120), e (-219) and f (-242). (D) mRNA fold change obtained targeting the endogenous gene NbAN2 with dCasEV2.1 through the multiplexing gRNA strategy after 4 dpi. Represented gRNAs target the promoter of NbAN2 in the positions: a (-103), b (-175), c (-196), d (-145), e (-198) and f: (-252). (E) mRNA fold change obtained targeting the endogenous gene NbDFR with dCasEV2.1 through the multiplexing gRNA strategy after 4 dpi. Represented gRNAs target the promoter of NbDFR at the positions: (-88), b (-125), c (-217), e (-198) and f (-248). In this experiment Myb-TF NbAN2 was included the, to test the RTA obtained with the natural transcription factor of NbDFR. (F) mRNA fold change obtained by the indirect activation of NbDFR at 7 dpi, by targeting NbAN1 and NbAN2 promoter genes with gRNAs a, b, c and d, e, f using dCasEV.2.1.

Next, we analysed the ability of dCasEV2.1 to induce endogenous *N. benthamiana* genes. For this purpose, we targeted the endogenous *N. benthamiana* homologue of DFR gene (NbDFR) and two transcription factors involved in polyphenol biosynthesis (the Myb factor NbAN2, and the bHLH protein NbAN1), which jointly activate NbDFR expression. In these experiments we followed a multiplexing strategy, targeting each gene with groups of three gRNAs expressed in multicistronic transcripts as depicted in Fig 3B. Results shows a strong 10000 X activation of NbDFR with dCasEV2.1 loaded with six gRNA combination, outperforming that obtained with endogenous Myb NbAN2 (Fig 3E). Similarly, NbAN1 and NbAN2 reached maximum (>1000X) activation rates with a 6X multiplex gRNA combination (Fig 3C, 3D). Simultaneous targeting of NbAN1 and NbAN2 led to an 80X secondary activation of NbDFR as shown in Fig 3F

The strong activations observed in endogenous genes when targeted with dCasEV2.1 prompted us to investigate if enhanced transcriptional activation had occurred at the expenses of specificity, therefore affecting unspecific or general changes in the transcriptome besides those on the designated target. To investigate this, we performed RNAseq analysis of leaf samples treated with dCasEV2.1 targeting the NbDFR and NbAN2 promoter. In parallel, leaf samples treated with dCasEV2.1 but no gRNAs were used as a control. The NbDFR-induced samples target NbDFR (Niben101Scf00305g05035) promoter with 6xgRNA (Fig 4A), following the gRNA design used in previous assays. Also, a search of potential off-targets was performed in order to find other possible genes activated in the transcriptome (Supplementary Table 8). A second gene (Niben101Scf00606g02015), labelled as NbDFR(h), known to be a closely related paralogue of selected NbDFR shares a 100% gRNA identical target in its promoter and a 2 mismatches homologue sequence (Fig 4B). The differential gene analysis is represented in Fig 4C and plots in the y axis the fold change between NbDFR-induced and control samples, whereas the x axis shows the log CPM indicative of absolute expression levels for each gene. As can be observed, the transcriptome of the dCasEV2.1/NbDFR sample, compared to the control, remains virtually unchanged with the exception of the NbDFR gene itself and a second gene, NbDFR(h). Strikingly, NbDFR is induced from extremely low expression levels to the top 50 mRNA in the transcriptome. No significant category enrichments of upregulated genes (Supplementary Data 1) and downregulated genes (Supplementary Data 2) were found in NbDFR-activated samples. The NbAN2-induced samples target NbAN2 (Niben101Scf00156g02004) promoter with 6gRNAs (Fig 4D) and a search of potential off-targets derived of this gRNAs was also performed (Supplementary table 9). An NbAN2 homologue (Niben101Scf00285g10004), labelled as NbAN2(h), also showing two 100% identical gRNA target sequences and three homologue sequence with at least 3 mismatches (Fig 4E). The differential gene analysis plotted in Fig 4F shows the equivalent representation obtained with dCasEV2.1 activation of NbAN2 gene. NbAN2 is now detected as the most activated gene in the genome. In this case, a second gene with a smaller fold change was observed, corresponding to NbAN2(h). Significant secondary activation of NbDFR was also detected at the transcriptome level, with much lower log CPM and fold change values. As expected for a TF, NbAN2 activation was accompanied by more significant changes in the transcriptome, although most of them had fold changes far below those observed in the targeted genes. GO analysis showed an enrichment in defence categories in upregulated genes (Supplementary Data 3), and primary metabolism and photosynthesis-related categories in downregulated genes (Supplementary Data 4).

**Figure 4.**
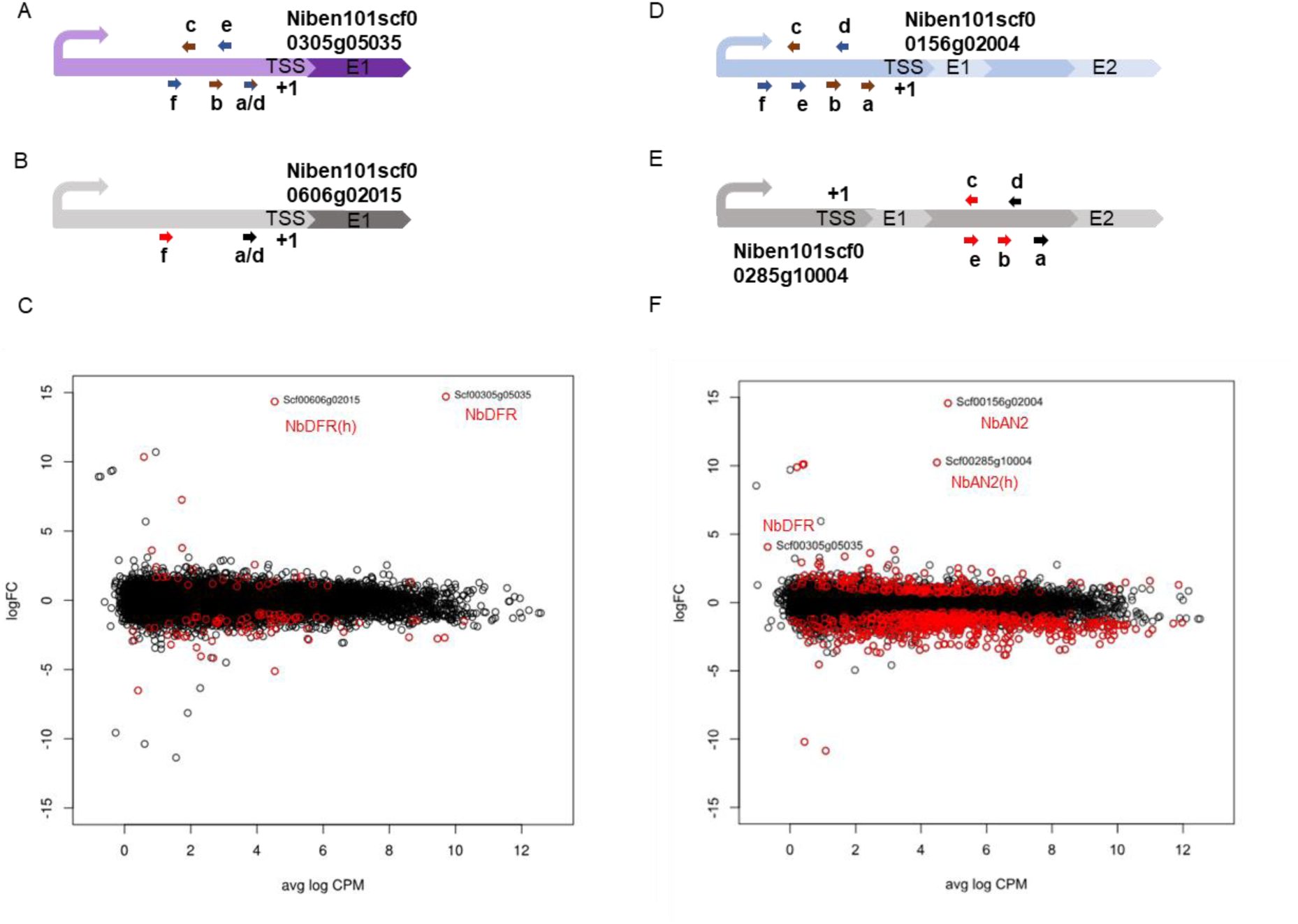
Differential gene expression analysis in samples activated with dCasEV2.1. (A) Representation of the NbDFR (Niben101scf00305g05035) promoter and exon 1 region with gRNAs targeted at positions: a/d (-88), b (-125), c (-217), e (-198) and f (-248) to TSS (B) Representation of homologous NbDFR (Niben101scf00606g02015) targeted at positions a/d (-89) and f (-261) to TSS. Black coloured gRNAs have 100% of identity in the homologous gene sequence, red coloured are off-targets gRNAs with at least 2 mismatches and off-target score <5. (C) Differential gene expression plot between the dCasEV2.1–NbDFR condition and control condition. The logFC axis represents the log of the fold change of the gene expression obtained in the samples that target the NbDFR with dCasEV2.1. Positive logFC values represents gene induction and negative logFC values represent gene repression. Avg logCPM is a normalized value for represent the gene abundance in the transcriptome. (D) Representation of the NbAN2 (Niben101scf00156g02004) promoter and exon 1 region with gRNAs targeted at positions: a (-103), b (-175), c (-196), d (-145), e (-198) and f: (-252) to TSS. (E) Representation of homologous NbAN2 (Niben101scf00285g10004) targeted at positions: a (+496), b (+424), c (+403), d (+474) and e (+401) to ATG. Black coloured arrows represent gRNAs with 100% of identity in the homologous gene sequence, red coloured arrows represent gRNAs off-targets with at least 3 mismatches and off-target score <2. (D) Differential gene expression plot between the dCasEV2.1–NbAN2 condition and control condition. The logFC axis represents the log of the fold change of the gene expression obtained in samples that target the NbAN2 with dCasEV2.1. Positive logFC values represents gene induction and negative logFC values represents gene repression. Avg logCPM is a normalized value for represents the gene abundance in the transcriptome.

## DISCUSSION

Ideally, programmable transcriptional activators should be potent, orthogonal, and able to activate a wide spectrum of targets. Maximum potency is an obvious engineering objective in the first place, as lower activation rates can be later obtained if needed by simply modifying the optimal parameters. The first generation of dCas9-PTAs tested in plants showed limited activation potency. Based on single translational fusions of dCas9 with activation domains, they have been soon overpassed by more sophisticated architectures involving tandem repeats of TAD and showing improved activation rates. Li et al ^21^ reported potent activation using the so-called TV combination, which comprised a VP128 tandem repeat with up to six copies of the TALE TAD motif (dCasTV strategy). This TV autonomous activator arranged in a SAM strategy (linked to the gRNA using the MS2 aptamer), produced strong activation rates in the OsGW7-LUC reporter construct assayed in rice protoplasts. Similarly, Lowder et al. ^16^ reported strong activation results with gRNA2.0 strategy, with maximum activation levels obtained with the so-called CRISPR-Act2.0 approach, which comprises a dCas9:VP64 coupled with MS2:VP64 via gRNA2.0 system.

In a search for further improvements in the performance of dCas9-PTAs, we designed, to our knowledge, the most comprehensive analysis of dCas9-PTAs tested in plants, comprising a total of 43 combinations of TADs displayed in different protein architectures, including those showing strongest activation in previous studies. When assayed against pNos:Luc reporter, the selected dCasEV2.1 PTA almost tripled the activation levels obtained with dCas9:TV and CRISPR-Act2.0 PTAs. Furthermore, a record activation of four orders of magnitude was reached when dCasEV2.1 targeted the endogenous NbDFR gene. The clue for this strong activation capacity seems to be an additive effect of the TADs employed (EDLL plus VPR) and the favourable gRNA2.1 loop. EDLL/VPR pairing was tested in the two possible combinations (dCas9:EDLL-MS2:VPR and vice versa), and in both of them the EDLL/VPR pair yielded highest activation rates as measured with pNos:Luc. On the other hand, the modification introduced by serendipity in the gRNA2.1 loop consistently yielded activation rates between 20-35% higher than the original gRNA2.0 loop in all different combination assayed. The causes of this effect are unknown, although we speculate that the modification could favour the stability of the gRNA scaffold in the plant cell. Besides dCasEV2.1, the remaining TAD combinations analysed here will be useful, alone or in combination with less efficient/distant gRNAs, to achieve intermediate activation levels when required. All the DNA elements showed here are integrated in the GoldenBraid 3.0 (GB3.0) modular cloning system and conform with the phytobrick standard^19,20^. GB3.0 enables fast combinatorial assembly, easy multigene engineering, exchange and reuse of standard parts and straightforward multiplexing of Cas9 gRNAs.

dCasEV2.1 successfully activated all promoter targets assayed, either imbibed in episomal T-DNAs or stably integrated in the chromosome driving expression of reporter constructs or endogenous genes, suggesting a wide activation spectrum that will need to be confirmed in further experiments involving a larger number of targets. However, the activation rates differed strongly among promoters. By referring all Fluc/Rluc measurements to those obtained with a standard pNos:Luc reporter phytobrick assayed in parallel, it was possible to obtain a general and comparative view of the transcriptional levels conferred to each of the three promoters assayed in transient expression analysis. A relationship between the basal expression levels of the promoter and the induction rates obtained with dCasEV2.1 became apparent by analysing these results, as it has been suggested earlier^22,23^. Thus, although the strongest induction rate was obtained with pSlDFR, this promoter could not reach levels above 5.9 RPUs in its fully activated state, which is half the activity measured for the CaMV35S-derived phytobrick. In contrast, pSlMTB, showing much more modest activation rates (8X), achieved instead remarkable RTA levels (80 RPUs), six times stronger than the 35S phytobrick. This was also confirmed in the stably-transformed reporter lines, with pSlMTB showing extremely high Fluc/Rluc levels upon induction, although in this case standard measurements are not possible due to the lack of a reference free of positional effects. The record Fluc/Rluc expression levels obtained with dCasEV2.1-activated pSlMTB suggest that this strategy could be exploited to boost yields of recombinant proteins and/or metabolites in biofactory and/or metabolic engineering approaches.

From an engineering standpoint, one of the most important features of PTAs is the ability to combine strong activation with genome-wide specificity. Off-target activities had been often regarded as the Achilles heel of Cas9 strategies, but their effect in plant PTA-regulation has not been established. Li et al. ^21^ showed almost no influence of dCas9:TV on the transcriptome profile of *Arabidopsis* protoplasts, however in that study the activation rates reached by the target gene were rather modest (below 10 fold). With RT-PCR data of DFR and AN genes showing activation rates between 1000-10000 folds, it was important to determine the effect on specificity. Deliberately, we selected NbDFR and NbAN2 target genes as representative of two distinct categories of potential actuators: (i) enzyme-coding genes (NbDFR) as final actuators with no transcriptional regulatory roles known, and (ii) transcriptional factors (NbAN2) representing a regulatory node with connections in the transcriptome. The results obtained with NbDFR-treated transcriptome demonstrate that strong activation with dCasEV2.1 is not incompatible genome-wide specificity, as an almost invariant profile was observed when compared with a control transcriptome, with the exception of an NbDFR homologue that served as additional proof of specificity. In contrast, dCasEV2.1-NbAN2 treatment resulted in wider transcriptome changes, generally consisting in low but significant changes in expression ratios. The changes observed in dCasEV2.1-NbAN2 are interpreted as a result of NbAN2 regulatory role. This is confirmed by the presence of NbDFR among the pool of genes significantly induced in dCasEV2.1-NbAN2 treatment, although with a fold change (and absolute expression levels) much lower than those observed in direct dCasEV2.1- NbDFR treatment. Interesting, dCasEV2.1-NbAN2 downregulated genes are enriched in GOs related with primary metabolism and photosynthesis, whereas upregulated genes are enriched in categories related with plant defence. It was earlier described that agroinfiltration of *N. benthamiana* Lab strain with Rosea-like Myb factors fails to engage a fully active anthocyanin biosynthesis pathway and initiates instead a necrosis-like programmed cell death process with activation of defensive pathways, a reaction fully compatible with the observed decrease in the expression of genes involved in primary metabolism ^24^.

In sum, we show here an improved PTA tool that combines maximum potency with genome-proven orthogonality. This tool has been shown to act efficiently at two different levels: with a heavily connected master regulator and with a final actuator enzyme. Both examples have strong potential applied implications. Targeting master regulators can be used to couple agronomically-relevant regulatory networks (e.g. defence, phase transition) to new external inputs, (e.g. agrochemicals); on the other hand, custom-activation of individual enzymatic steps will facilitate steering metabolic flows towards selected final products.

## METHODS

### GB phytobricks construction and assembly

Llevel 0 used in this work were created following the domestication strategy described in Sarrion-Perdigones et al. ^25^. Plasmids pHRdSV40-dCas9-24xGCN4-v4-P2A-BFP, Addgene ID: #60903 and pHRdSV40-scFv-GCN4-sfGFP-VP64-GB1-NLS, Addgene ID #60904 ^12^ kindly provided by Ron Vale laboratory, served as a template for the construction of GB2464 and GB1463 by PCR amplification, using the Phusion High Fidelity DNA polymerase. All level 0 parts are listed in Supplementary Table 1 and their sequences can be searched at https://gbcloning.upv.es with the corresponding GB IDs. Multipartite BsaI restriction–ligation reactions from level 0 parts and binary BsaI or BsmBI restriction–ligation reactions were performed as described in ^25^ to obtain all the level ≥1 assemblies. A list with all the TUs and modules used in this work is provided on Supplementary Table 5. All level ≥1 constructs were validated by restriction enzyme analysis.

All gRNAs used in this work were designed using the Benchling CRISPR tool (https://benchling.com) and following the schema described in Supplementary Figure 1A, for gRNA position determination.

For single gRNA assembly in GB level 1 (Supplementary Figure 1B), primers including the protospacer sequence were designed at http://www.gbcloning.upv.es/do/crispr/. All primers used for gRNAs assembly are listed in Supplementary Table 6. Primers were resuspended in water to final concentrations of 10 µM. Equal volumes of forward and reverse primers for each gRNA were mixed. The mixture was incubated at room temperature for 5 min for the hybridization of the primer pair. gRNA assembly in level 1 was carried out with a BsaI restriction–ligation reaction. The reactions were set up in 10 µl with 1 µl of primers mix, 75 ng of U626 promoter (GB1001), 75 ng of the corresponding scaffold (GB0645, GB1436, GB2461, GB1437, GB1450 or GB1451) and 75 ng of pDGB3α destination vector. All gRNAs in level 1 used in this work are listed in Supplementary Table 4.

For the assembly of gRNAs to be used in the multiplexing strategy, GB level -1 plasmids containing the tRNA and the gRNA2.1 scaffold were designed following the plasmid structure described in Vazquez-Vilar et al.^10^. For the construction of these plasmids, DNA fragments including the tRNA and the gRNA2.1 scaffold were synthesized as IDT gBlocks® Gene Fragments and subsequently cloned in pVD1 (GB0101) with a BsaI restriction-ligation reaction. Level -1 plasmids are listed in Supplementary Table 2. Individual gRNAs were assembled in pUPD2 with a BsmBI restriction-ligation reaction that was performed with 75 ng of pUPD2 and 75 ng of the corresponding level –1 pVD1; pVD1_M1-3pTRNA scf 2.1, pVD1_M2-3pTRNA scf 2.1 or pVD1_M3-3pTRNA scf 2.1 plasmid, depending on the desired position of each target on the polycistronic gRNA and a mix of complementary primers with the protospacer sequence. Supplementary Figure 1C shows the schematic representation of the multiplexing gRNAs cloning strategy. All level 0 gRNAs generated with this cloning strategy are listed in Supplementary Table 3. All constructs were validated by RE-analysis and confirmed by sequencing.

### *Nicotiana benthamiana* agroinfiltration

The transient expression assays were carried out through agroinfiltration of *N. benthamiana* leaves. *N. benthamiana* plants were grown for 5 weeks before agroinfiltration in a growing chamber where the growing conditions were 24°C/20°C light/darkness with a 16h/8h photoperiod. The plasmids were transferred to *Agrobacterium tumefaciens* strain GV3101 by electroporation. Agroinfiltration was carried out with overnight grown bacterial cultures. The cultures were pelleted and resuspended on agroinfiltration solution (10 mM MES, pH 5.6, 10 mM MgCl2, and 200 μM acetosyringone). After incubation for 2 h at room temperature with agitation, the optical density of the bacterial cultures was adjusted to of 0.1 at 600nm. Cultures were mixed in equal volumes for co-infiltration. The silencing suppressor P19 was included in all the assays; in the same T-DNA for the transcriptional regulation experiments with the reporter constructs and co-delivered in an independent T-DNA for the transcriptional regulation of endogenous genes assays. Agroinfiltrations were carried out through the abaxial surface of the three youngest fully expanded leaves of each plant with a 1 ml needle-free syringe.

### Luciferase/Renilla activity determination

The assays conditions follow the experimental standards found in https://gbcloning.upv.es/add/experiment/SE_002 with minor modifications. Samples were collected at 5 days post infiltration (dpi) instead of 4 dpi. For the determination of the luciferase/renilla activity one disc per leaf (d = 0.8 cm, approximately 18–19 mg) was excised, homogenized and extracted with 375µl of ‘Passive Lysis Buffer,’ followed by 10 min of centrifugation (14,000×g) at 4 °C. Then, the supernatant was collected as working plant extract. Fluc and Rluc activities were determined following the Dual-Glo® Luciferase Assay System (Promega) manufacturer’s protocol with minor modifications:

7.5 µl of working plant extract, 30 µl of LARII and 30 µl of Stop&Glow Reagent were used. Measurements were made using a GloMax 96 Microplate Luminometer (Promega) with a 2-s delay and a 10-s measurement. Fluc/Rluc ratios (RPUs) were determined as the mean value of three biological replicates coming from three independent agroinfiltrated leaves of the same plant and were normalized to the Fluc/Rluc ratio obtained for a reference sample, that measures the basal activity of the evaluated promoter fused to the reporter.

### Generation and selection of *N. benthamiana* reporter lines.

The *N. benthamiana* reporter pNos, SlDFR and SlMTB lines were generated following the transformation protocol described previously ^26^. Constructs GB2248 (pNos), GB2250 (SlDFR), GB2249 (pMTB) were transferred to LBA4404 *Agrobacterium tumefaciens* strain use for plant transformation. Murashige and Skoog (MS) plates supplied with Kanamycin at 100 mg/ml were used to select the transgenic T0 lines. T1 plants were selected for single copy T-DNA insertions based on the segregation of their offspring. Single-copy T2 lines were sorted for further analysis on the basis of their luciferase/renilla activity rates. For pNos, three homozygous T2 lines representing low (Nos-RL6), medium (Nos-RL3) and high (Nos-RL5) Fluc/Rluc expression levels were selected. For SlMTB, analysis was conducted using the T2 homozygous line showing higher expression levels (MTB-RL3). Finally, T2 heterozygous DFR-RL1 reporter line was selected on the basis of its high Fluc/Rluc induction rates when agroinfiltrated with a Ros1 TF. All selected lines were grown under 24°C/20°C light/darkness with a 16h/8h photoperiod conditions in a growing chamber.

### RNA isolation and qRT-PCR Gene Expression Analysis.

Total RNA was isolated from 100mg of fresh leaf tissue harvested 4 and 7 dpi using RNA isolation-kit Macherey-Nagel according to the manufacturer’s instructions. Before cDNA synthesis, total RNA was treated with rDNAse-I Invitrogen Kit according to the manufacturer. An aliquot of 1µg of total RNA was used for cDNA synthesis using PrimeScript™ RT-PCR Kit (Takara) in 20 µL final volume according to the manufacturer. Expression levels of each gene were measured in triplicated reactions, performed with the same cDNA pool of each condition, in presence of florescent dye (SYBR® Premix Ex Taq) using Applied biosystem 7500 Fast Real Time PCR system with specific primer pairs (Table 7, supplemental). F-BOX protein was used as internal reference gene^27^. Basal expression levels were calculated either with samples agroinfiltrated with P19 or samples with the dCas9-activation TUs without gRNAs. Calculations of each sample were carried out according the comparative ΔΔCT method^28^.

### RNA sequencing and analysis.

RNA samples were collected and isolated following the protocol for RNA isolation described above. Three biological replicates were selected for each condition; control condition, NbDFR activation and NbAN2 activation. The control condition was agroinfiltrated with the T-DNA that contains the dCasEV2.1 construct without any gRNA. RNA-sequencing was undertaken using TruSeq Stranded mRNA LT Sample Prep Kit for library construction. Libraries were sequenced using Illumina Hiseq 4000 platform. 40 million 150bp paired-end reads per sample were generated and and mapped using *N. benthamiana* genome Niben v1.0.1 as reference, available at https://solgenomics.net/, through Hisat2 programm ^29^. The quality of the reads obtained was evaluated with FastQC programm available online at: http://www.bioinformatics.babraham.ac.uk/projects/fastqc. Furthermore, the Illumina adaptors were eliminated from the reads using the program Trimmomatic ^30^. For each sample a count of the expression was performed using the programm StringTie^31^, following the gene models for the genome reference Niben v1.0.1, also available in https://solgenomics.net/. A statistical analysis with the raw data was performed in order to evaluate the differential expression between the samples NbDFR and NbAN2 against control samples, using a statistical package edgeR ^32^. For each comparison of the expression data, the results were organized depending on the profile of the expression based on the up-regulation or down-regulation of the genes and GO term analysis was also performed using Blast2GO ^33^ and using FDR<0.05. The off-targets analysis was performed through Benchling (https://benchling.com) and Cas-OFFinder ^34^ allowing until 4 nucleotides of mismatch. Possible off-targets located within 1000bp upstream and 500bp down-stream the TSS of the genes were analysed to see if they were differentially expressed ^35^.

The reads have been deposited in the Sequence Read Archive (SRA), under the Bioproject PRJNA507084 (http://www.ncbi.nlm.nih.gov/bioproject/507084)

## Supporting information

## ACKNOWLEDGMENTS

This work has been funded by Grant BES-2017-080098 from Plan Nacional I+D of the Spanish Ministry of Economy and Competitiveness. Selma S. is a recipient of FPI fellowship associated to this Grant. Bernabé-Orts J is a recipient of FPI associated to Grant BIO2013-42193 Plan Nacional I+D of the Spanish Ministry of Economy and Competitiveness. The authors would like to thank Asun Fernandez del Carmen for assistance with the manuscript preparation.

