## Supplementary material for "Strong gene activation with genome-wide specificity using a new orthogonal CRISPR/Cas9-based Programmable Transcriptional Activator"

**Table 1**: GB level 0 parts used and generated available in <http://www.gbcloning.upv.es/>

| **Nº GB** | **Name** | **GB Nickname** |
| --- | --- | --- |
| **GB1079** | pUPD2:dCas9 | pdCas9 |
| **GB1001** | pUPD: U6-26 | pU6-26 |
| **GB0645** | pUPD2: sgRNA | psgRNA |
| **GB1436** | pUPD2: sgRNA aptamer SAM | pUPD2 sgRNA:scaffold tetraloop MS2 aptamer |
| **GB2461** | pUPD2:sgRNA2.0 scRNA | pUPD2_sgRNA sCRNA 2.0 |
| **GB1437** | pUPD2: sgRNA2.1 scRNA | pUPD2_sgRNAscaffold 2xF6Ms2 Aptamer |
| **GB1450** | pUPD2: sgRNA PP7 aptamer | pUPD2 PP7 Stem Loop |
| **GB1451** | pUPD2: sgRNA COM aptamer | pUPD2 scf_COM |
| **GB2464** | pUPD: SunTag | SunTag |
| **GB1463** | pUPD2: ScFv | pUPD2_ScFV |
| **GB1435** | pUPD2:MS2 | pUPD2_MS2 |
| **GB1453** | pUPD2:PP7 | pUPD2 NLS-PP7 |
| **GB1786** | pUPD2:COM | pUPD2:NLS-COM1 |
| **GB1186** | pUPD2:VP64 | p3xNLS-VP64 |
| **GB1187** | pUPD2: EDLL | p3xNLS-EDLL |
| **GB1791** | pUPD2: p300 core | pUPD2_P300core |
| **GB1850** | pUPD2: VP192 | pUPD2_VP192 |
| **GB1814** | pUPD2: VPR | pUPD2_VPR |
| **GB1817** | pUPD2:ERF2(M) | pUPD2_ERF2 |
| **GB2001** | pUPD2:TV | pUPD2_TV |
| **GB0030** | pUPD: p35s | pP35S |
| **GB0037** | pUPD:tNos | pTnos |
| **GB2382** | pUPD2:NbAN2 | pUPD2_NbAN2 |

**Table 2:** GB level -1 parts for multiplexing strategy

| **Nº GB** | **Name** | **GB database Nickname** |
| --- | --- | --- |
| **GB2073** | pVD1_M1-3pTRNA scf 2.1 | pVD1_M1-F6-pTRNA-scf |
| **GB2074** | pVD1_M2-3pTRNA scf 2.1 | pVD1_M2-F6-pTRNA-scf |
| **GB2075** | pVD1_M3-3pTRNA scf 2.1 | pVD1_M3-F6-pTRNA-scf |

**Table 3**: GB gRNAs level 0 for multiplexing assembly

| **Nº GB** | **Name** | **GB database Nickname** |
| --- | --- | --- |
| **GB2148** | Multiplex M1_3gRNA-88 NbDFR 2.1 | pUPD2_M1-F6-sgRNANbDFR-85 |
| **GB2149** | Multiplex M2_3gRNA-125 NbDFR 2.1 | pUPD2_M2-F6-sgRNANbDFR-145 |
| **GB2150** | Multiplex M2_3gRNA-198 NbDFR 2.1 | pUPD2_M2-F6-sgRNANbDFR-198 |
| **GB2151** | Multiplex M3_3gRNA-217 NbDFR 2.1 | pUPD2_M3-F6-sgRNANbDFR-218 |
| **GB2152** | Multiplex M3_3gRNA-248 NbDFR 2.1 | pUPD2_M3-F6-sgRNANbDFR-268 |
| **GB2153** | Multiplex M1_3gRNA-103 NbAN2 2.1 | pUPD2_M1-F6-sgRNANbAN2-103 |
| **GB2154** | Multiplex M1_3gRNA-125 NbAN2 2.1 | pUPD2_M1-F6-sgRNANbAN2-125 |
| **GB2155** | Multiplex M2_3gRNA-175 NbAN2 2.1 | pUPD2_M2-F6-sgRNANbAN2-175 |
| **GB2156** | Multiplex M2_3gRNA-198 NbAN2 2.1 | pUPD2_M2-F6-sgRNANbAN2-198 |
| **GB2157** | Multiplex M3_3gRNA-196 NbAN2 2.1 | pUPD2_M3-F6-sgRNANbAN2-196 |
| **GB2158** | Multiplex M3_3gRNA-252 NbAN2 2.1 | pUPD2_M3-F6-sgRNANbAN2-252 |
| **GB2302** | Multiplex M1_3gRNA-101 NbAN1 2.1 | pUPD2_M1-3NbAN1_4283-101 |
| **GB2303** | Multiplex M1_3gRNA-120 NbAN1 2.1 | pUPD2_M1-3NbAN1_4283-120 |
| **GB2304** | Multiplex M2_3gRNA-173 NbAN1 2.1 | pUPD2_M2-3NbAN1_4283-173 |
| **GB2305** | Multiplex M2_3gRNA-219NbAN1 2.1 | pUPD2_M2-3NbAN1_4283-219 |
| **GB2306** | Multiplex M3_3gRNA-242 NbAN1 2.1 | pUPD2_M3-3NbAN1_4283-242 |
| **GB2307** | Multiplex M3_3gRNA-261 NbAN1 2.1 | pUPD2_M1-3NbAN1_4283-261 |

**Table 4**: GB gRNA Level 1. Standard and Multiplexing strategy

| **Nº GB** | **Name** | **GB database Nickname** |
| --- | --- | --- |
| **GB1197** | U6-26-gRNA-161 pNos | pEGB U626:gRNA4pNOS:sgRNA |
| **GB2462** | U6-26-gRNA-161 pNos scf 2.0 | 3alpha1_U6-26-gRNA4pNos scf aptamer 2.0 Native |
| **GB1724** | U6-26-gRNA-161 pNos scf 2.1 | pDGB3_alpha1_U6-26:gRN A 4 (pNos):MS2 F6x2 aptamer |
| **GB1725** | U6-26-gRNA-161 pNos scf SAM | pDGB3_alpha1_U6-26:gRNA 4 (pNos):MS2 tetraloop and loop aptamer |
| **GB1740** | U6-26-gRNA-161 pNos scf PP7 aptamer | pDGB3_alpha1_U6-26:gRNA4(pNos):PP7stem loop |
| **GB1797** | U6-26-gRNA-161 pNos scf COM aptamer | 3alpha1_U6-26-gRNA4pNos COM scf |
| **GB1744** | U6-26-gRNA-211 pNos scf 2.0 * | pDGB3_alpha1_U6-26:gRNA5(pNos):MS2 F6x2 aptamer |
| **GB2049** | U6-26-gRNA -161Pnos scf 2.1 -35s:dCas9:EDLL:Tnos-U6-26-gRNA-211 Pnos scf 2.0* - 35s:MCP:VPR:Tnos | pDGB3_alpha2_U6-26-4gRNA Pnos-F6x2_35s-dCas9:EDLL-Tnos-U6-26-5gRNA Pnos scf F6x2 - 35s-Ms2:VPR-Tnos |
| **GB2045** | U6-26-gRNA-98 SlMTB scf 2.1 | 3alpha1:U6-26-2gRNA_MTB_scf F6x2 |
| **GB1801** | U6-26-gRNA-129 SlMTB scf 2.1 | 3alpha1_U6-26_MTB3-sgRNa-MS2 F6x2 scf |
| **GB2044** | U6-26-gRNA-184 SlMTB scf 2.1 | 3alpha2:U6-26-4gRNA_MTB_scf F6x2 |
| **GB1859** | U6-26-gRNA-541 SlMTB scf 2.1 | 3alpha1:U6-26-5gRNA_MTB_scf F6x2 |
| **GB2070** | U6-26-gRNA-541 SlMTB scf 2.1 + U6-26-gRNA-541 SlMTB scf 2.1 + U6-26-gRNA-129 SlMTB scf 2.1 + U6-26-gRNA-98 SlMTB scf 2.1 | pDGB3_Alpha1_U6-26-5gRNA MTB-F6x2_U6-26-4gRNA MTB-F6x2_U6-26-3gRNA MTB-F6x2_U6-26-2gRNA MTB-F6x2 |
| **GB1838** | U6-26-gRNA-150 SlDFR scf 2.1 | 3alpha1_U6-26-1gRNA-DFR F6x2 MS2scf |
| **GB1837** | U6-26-gRNA-300 SlDFR scf 2.1 | 3alpha1_U6-26-4gRNA-DFR F6x2 MS2scf |
| **GB1839** | U6-26-gRNA-376 SlDFR scf 2.1 | 3alpha1_U6-26-5gRNA-DFR F6x2 MS2scf |
| **GB2169** | Multiplex scRNA 2.1 U6-26 gRNA -88, -125,-217 NbDFR | 3Alpha1_U6-26-sgRNANbDFR -85,-145,-218 F6x2 Multiplex |
| **GB2170** | Multiplex scRNA 2.1 U6-26 gRNA -88, -198,-248 NbDFR | 3Alpha2_U6-26-sgRNANbDFR -85,-198,-268 F6x2 Multiplex |
| **GB2176** | Multiplex scRNA 2.1 U6-26 gRNA -88, -125,-217 NbDFR + Multiplex scRNA 2.0 U6-26 gRNA -88, -198,-248 NbDFR | 3Omega2_U6-26-sgRNANbDFR -85,-145,-218 + U6-26-sgRNANbDFR -85,-198,-268 F6x2 Multiplex |
| **GB2171** | Multiplex scRNA 2.1 U6-26 gRNA -103, -175,-196 NbAN2 | 3Alpha1_U6-26-sgRNANbAN2 -103,-175,-196 F6x2 Multiplex |
| **GB2172** | Multiplex scRNA 2.1 U6-26 gRNA -125, -198,-252 NbAN2 | 3Alpha2_U6-26-sgRNANbAN2 -125,-198,-252 F6x2 |
| **GB2177** | Multiplex scRNA 2.1 U6-26 gRNA -103, -175,-196 NbAN2 + Multiplex scRNA 2.0 U6-26 gRNA -125, -198,-252 NbAN2 | 3Omega2_U6-26-sgRNANbAN2 -103,-175,-196 + U6-26-sgRNANbAN2 -125,-198,-252 F6x2 |
| **GB2309** | Multiplex scRNA 2.1 U6-26 gRNA -101, -173,-242 NbAN1 | 3alpha 1 Multiplex U626_NbAN14283-101-173-242 |
| **GB2310** | Multiplex scRNA 2.1 U6-26 gRNA -120, -219,-261 NbAN1 | 3alpha 2 Multiplex U626_NbAN14283-120-219-261 |
| **GB2408** | Multiplex scRNA 2.1 U6-26 gRNA -101, -173,-242 NbAN1 + Multiplex scRNA 2.0 U6-26 gRNA -120, -219,-261 NbAN1 | Omega2:U2626_NbAN1_4283_-101-173-242 F6 + U2626_NbAN1_4283_-120-219-261 F6 |

**Table 5:** GB level 1 and >1 TUs and Modules

| **Nº GB** | **Name** | **GB database Nickanme** |
| --- | --- | --- |
| **GB1398** | pNos:Luciferase:tNos-35S:Renilla:tNos-35S:P19:tNos | pEGB3alpha2 Pnos:luc:Tnos-SF-35S:Ren:Tnos-35s:P19:Tnos-SF |
| **GB2248** | tNos:NptII:Pnos-Pnos:Luc:tNos-35s:Ren:tNos | Tnos:NptII:Pnos-Pnos:Luc:Tnos-35s:Ren:Tnos |
| **GB1399** | SlMTB:luc:tNos-SF-35S:Ren:tNos-35s:P19:tNos-SF | pEGB3alpha2 MTB:luc:Tnos-SF-35S:Ren:Tnos-35s:P19:Tnos-SF |
| **GB2250** | tNos:NptII:Pnos-SIDFR:Luc:tNos-35s:Ren:tNos | Tnos:NptII:Pnos-SIDFR:Luc:Tnos-35s:Ren:Tnos |
| **GB1160** | SlDFR:Luc:tNos-SF-35S:Renilla:tNos-35S:P19:tNos | pEGB SlDFR:Luc:TNos-SF-35S:Renilla:TNos-35S:P19:TNos |
| **GB2249** | tNos:NptII:pNos-SlMTB:Luc:tNos-35s:Ren:tNos | Tnos:NptII:Pnos-MTB:Luc:Tnos-35s:Ren:Tnos |
| **GB0164** | p35s:Luciferase:tNos-35S:Renilla:tNos-35S:P19:tNos | 35s:Luciferase:Tnos-SF-35s:Renilla:Tnos-35s:P19:Tnos |
| **GB1603** | 35s:dCas9:SunTag:tNos | 35s:dCas9-SunTag:Tnos |
| **GB1189** | 35s:dCas9:VP64:tNos | pEGB 35s:dCas9-VP64:tNOS |
| **GB1190** | 35s:dCas9:EDLL:tNos | pEGB 35s:dCas9-EDLL:tNOS |
| **GB1794** | 35s:dCas9:p300:tNos | 3alpha2_35s-dCas9:P300_Tnos |
| **GB1851** | 35s:dCas9:VP192:tNos | 3alpha2_35s-dCas9:VP192-Tnos |
| **GB1826** | 35s:dCas9:VPR:tNos | 3alpha2_35s-dCas9:VPR-Tnos |
| **GB1824** | 35s:dCas9:ERF2:tNos | 3alpha2_35s-dCas9:ERF2-Tnos |
| **GB2047** | 35s:dCas9:TV:tNos | 3alpha2_35s-dCas9:TV-Tnos |
| **GB1403** | 35S:MS2-VP64:tNos | pDGB3alpha2 35S:MS2-VP64:Tnos |
| **GB1738** | 35s:MS2:EDLL:tNos | pDGB3_alpha2: 35s_MS2:EDLL_Tnos |
| **GB1852** | 35s-Ms2:VP192_tNos | 3alpha2_35s-Ms2:VP192_Tnos |
| **GB1830** | 35s:Ms2:VPR:tNos | 3alpha1_35s-MS2:VPR-Tnos |
| **GB1833** | 35s-MS2:ERF2-tNos | 3alpha2_35s-MS2:ERF2-Tnos |
| **GB2048** | 35s:MS2:TV:tNos | 3alpha2_35s-MS2:TV-Tnos |
| **GB1460** | 35S:PP7:VP64:tNos | pEGB3alpha1 35S:NLS-PP7-VP64:Tnos |
| **GB1476** | 35S:COM:VP64:tNos | 35s:COM-VP64:Tnos |
| **GB1592** | 35s:ScFv:VP64:tNos | 35S:scFv-VP64:Tnos |
| **GB1836** | 35s:ScFv:EDLL:tNos | 3alpha2_35s-ScFv-EDLL-tNOS |
| **GB2085** | 35s:Ms2:VPR:tNos-35s:dCas9:EDLL:tNos | pDGB3_Omega1_35s-Ms2:VPR-Tnos-35s-dCas9:EDLL-Tnos |
| **GB2384** | 35s:NbAN2:tNos | 3alpha2_35s:NbAN2:Tnos |
| **GB0127** | 35s:SlANT1:tNos | pEGB 35S:Ant1:Tnos |

**Table 6**: Protospacer sequence and position design for each promoter.

| **Promoter** | **Position to TSS** | **Strand** | **Sequence** |
| --- | --- | --- | --- |
| pNos | -161 | Coding | GCCACTGAGCCGCGGGTTTC |
| pNos | -211 | Coding | GGGACAAGCCGTTTTACGTT |
| SlMTB | -98 | Coding | TACGATCACGACACGTGTAC |
| SlMTB | -129 | Coding | GATGAAATTAGGATCATGTA |
| SlMTB | -184 | Coding | GTCTAGAACATACGTACGAA |
| SlMTB | -541 | Non coding | GATGTAGCATATGAGATGAT |
| SlDFR | -150 | Non coding | GACTGGTTGGTGAGAGAAGA |
| SlDFR | -300 | Non coding | GGATAAAATGGTAATAGTTT |
| SlDFR | -376 | Coding | GCTGTATCTAATAGAATCTT |
| NbDFR | -88 | Coding | ATGACTGACTGGTTGGTGAG |
| NbDFR | -125 | Non coding | TCCATATATAGATAAGAAAG |
| NbDFR | -198 | Coding | TATCCGTATGCCTTACCTTT |
| NbDFR | -217 | Coding | TTGGATTTTGGTGTATTCTT |
| NbDFR | -248 | Non coding | TTGAGAATTTGGTAAAACGA |
| NbAN2 | -103 | Coding | GGAGTTACGCTAATCACTAG |
| NbAN2 | -145 | Non coding | CAGTGCAATTTATTACTCAT |
| NbAN2 | -175 | Coding | CGTAAAAAGTCCATATCGAC |
| NbAN2 | -198 | Coding | GACGCGTAGCTCTCTCCAAT |
| NbAN2 | -196 | Non coding | GAACAGTGTCTACTGCCAAT |
| NbAN2 | -252 | Coding | TGTCCTTTTCACTATTAAGT |
| NbAN1 | -101 | Non coding | TAGGAGGAATGAGTGTGCGT |
| NbAN1 | -120 | Coding | AATAATAGTATAATAACCAA |
| NbAN1 | -173 | Coding | TTATACCGTAAGTAACTTAG |
| NbAN1 | -219 | Non coding | GTATTACGCTTTTAATCACT |
| NbAN1 | -242 | Non coding | CACAACAATCTAATTAAAAA |
| NbAN1 | -261 | Coding | ACCCGGGTCAAGCCGTGTAA |

**Table 7**: Primer pair for qRT analysis.

| **Target Gene** | **Sequence 5’-3’** |
| --- | --- |
| NbDFR | TTCATCTGCGCATCCCATCA |
|  | TCCCTACTGAGTTTAAAGGTATCGA |
| NbAN1 | CATCTCTTAATAATGGCGTCTTCTTG |
|  | CTCTAGGGATTATCTGATGTATTGACC |
| NbAN2 | GGAAAAGTTGCAGACTGAGGTG |
|  | ACCCGCAATAAGTGACCATCTG |
| NbF-box | TTGGAAACTCTCTCCCCACTTG |
|  | GCTCATTGTTGGATGGGTACCT |

**Table 8.** Potential off-targets of gRNAs used for NbDFR dCas9EV2.1 activation.

| **gRNA**  **position** | **Targeted Gene** | **Potential offtargets** | **Sign NbDFR-Control gene differential expression analysis.** |
| --- | --- | --- | --- |
| NbDFR -88 | Niben101Scf00305g05035 | Niben101Scf00311g05015(UP) Niben101Scf00606g02015(UP) Niben101Scf05634g02005(UP) | Niben101Scf00606g02015 |
| NbDFR -125 | Niben101Scf00305g05035 | Niben101Scf01792g00022(UP) Niben101Scf02399g01012(DOWN) Niben101Scf02764g05013(UP) Niben101Scf02764g05013(DOWN) Niben101Scf03455g04002(DOWN) Niben101Scf05217g01001(DOWN) Niben101Scf14642g04009(DOWN) | NONE |
| NbDFR -198 | Niben101Scf00305g05035 | Niben101Scf01607g06019(UP) Niben101Scf21739g00001(UP) | NONE |
| NbDFR -218 | Niben101Scf00305g05035 | Niben101Scf00320g02010(UP) Niben101Scf00320g02006(UP) |  |
| NbDFR -248 | Niben101Scf00305g05035 | Niben101Scf00606g02015(UP) | Niben101Scf00606g02015 |

**Table 9:** Potential off-targets of gRNAs used for NbAN2 dCas9EV2.1 activation

| **gRNA**  **position** | **Targeted Gene** | **Potential offtargets** | **Sign NbAN2-Control gene differential expression analysis** |
| --- | --- | --- | --- |
| NbAN2 -103 | Niben101Scf00156g02004 | Niben101Scf00285g10004(DOWN) | Niben101Scf00285g10004 |
| NbAN2 -145 | Niben101Scf00156g02004 | Niben101Scf00152g10003(UP) Niben101Scf00156g03002(UP) Niben101Scf00285g10004(DOWN) Niben101Scf02868g03005(UP) Niben101Scf03306g00002(DOWN) Niben101Scf08512g00009(DOWN) | Niben101Scf00285g10004 |
| NbAN2 -175 | Niben101Scf00156g02004 | Niben101Scf00285g10004(DOWN) Niben101Scf00288g15003(DOWN) Niben101Scf03414g03001(UP) Niben101Scf03518g00006(UP) | Niben101Scf00285g10004 |
| NbAN2 -198 | Niben101Scf00156g02004 | Niben101Scf00285g10004(DOWN) | Niben101Scf00285g10004 |
| NbAN2 -196 | Niben101Scf00156g02004 | Niben101Scf00285g10004(DOWN) | Niben101Scf00285g10004 |
| NbAN2 -252 | Niben101Scf00156g02004 | Niben101Scf01383g09023(UP) Niben101Scf05342g05013(UP) Niben101Scf14009g01001(UP) | NONE |

**Supplementary figure 1**

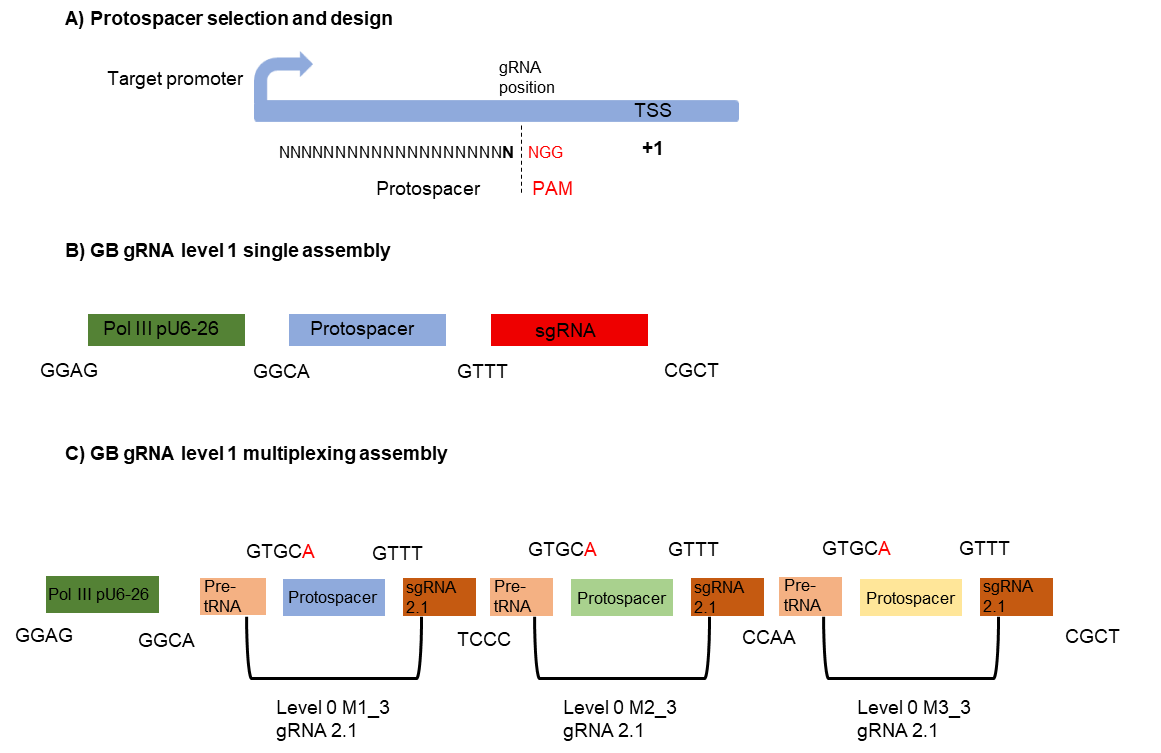

**Supplementary Figure 1: gRNA design and assembly**

(A) Protospacer selection for gRNA design. The gRNA position was established taking as a reference the Transcriptional Start Side (TSS) of the target promoter and the first nucleotide after the PAM sequence. (B) Representation of the single gRNA GB assembly. (C) Representation of the Multiplexing strategy for gRNA GB assembly with the gRNA scaffolds 2.1.

**Supplementary figure 2**

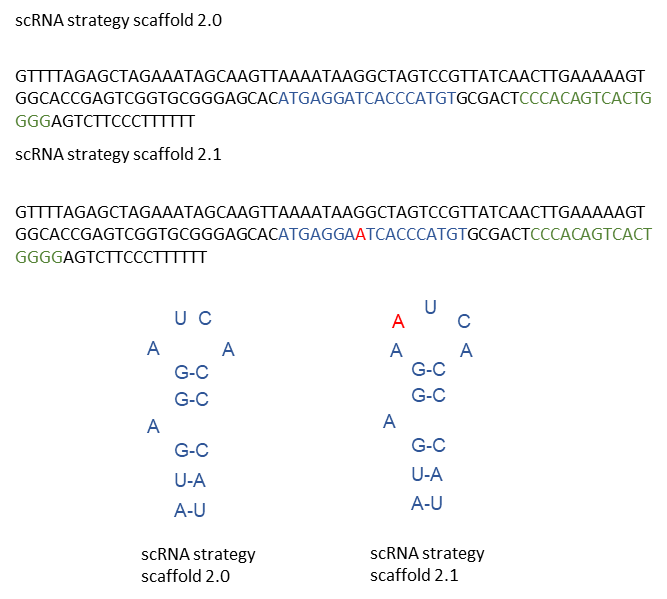

**Supplementary figure 2: ScRNA scaffold 2.0 and 2.1 representation**

Sequence of the scaffold 2.0 and scaffold 2.1 of the scRNA strategy. Representation of the adenine incorporation in MS2 loop aptamer of the scaffold 2.1 variant sequence.

**Supplementary figure 3**

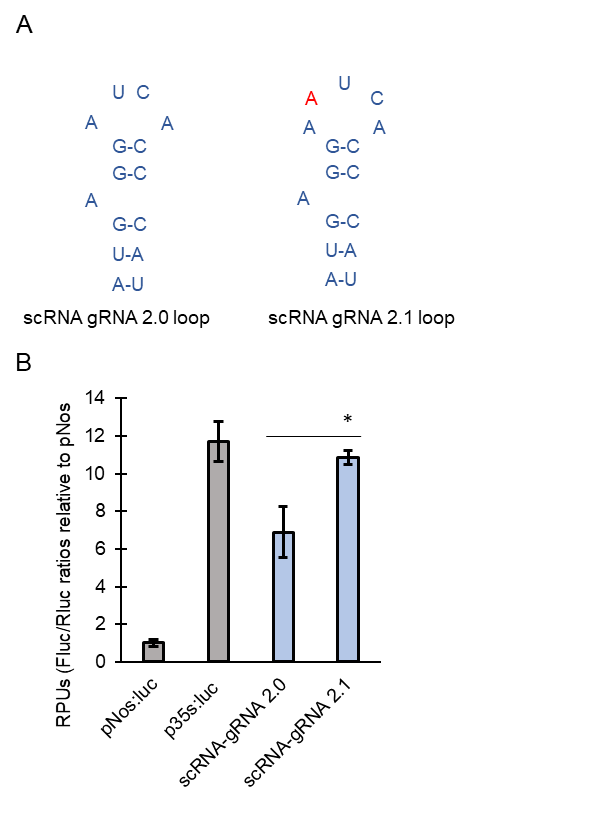

**Supplementary figure 3**: **Comparison of scRNA gRNA 2.0 and 2.1 using dCas9:EDLL-MS2:VPR targeting the pNos promoter.**

(A) Representation of the MS2 aptamer loop in 2.0 and 2.1 variant. (B) Relative transcriptional activities (RTAs) obtained with the gRNA 2.0 and 2.1 variant targeting pNos promoter at position -161 with dCas9EDLL-MS2:VPR. Asterisk represents T student significant values *p<0,05
